# Next generation CRISPR/Cas9 transcriptional activation in *Drosophila* using flySAM

**DOI:** 10.1101/252031

**Authors:** Yu Jia, Rong-Gang Xu, Xingjie Ren, Ben Ewen-Campen, Rajendhran Rajakumar, Jonathan Zirin, Donghui Yang-Zhou, Ruibao Zhu, Fang Wang, Decai Mao, Ping Peng, Huan-Huan Qiao, Xia Wang, Lu-Ping Liu, Bowen Xu, Jun-Yuan Ji, Qingfei Liu, Jin Sun, Norbert Perrimon, Jian-Quan Ni

**Affiliations:** Gene Regulatory Lab, School of Medicine, Tsinghua University, Beijing 100084, China; Institute for Human Genetics and Department of Neurology, University of California San Francisco, San Francisco, California 94143, USA; Department of Genetics, Harvard Medical School, Boston, MA 02115; Drosophila RNAi Screening Center, Department of Genetics, Harvard Medical School, Boston, MA 02115; Sichuan Academy of Grassland Science, Chengdu 611731, China; Tsinghua Fly Center, Tsinghua University, Beijing 100084, China; Howard Hughes Medical Institute, Boston, MA 02115; Department of Molecular and Cellular Medicine, College of Medicine, Texas A&M Health Science Center, College Station, TX 77843; School of Pharmaceutical Sciences, Tsinghua University, Beijing 100084, China; Tsinghua University-Peking University Joint Center for Life Sciences, Beijing 100084, China

**Author notes:** Y.J., R-G.X., X.R. and B.E-C contributed equally to this work.

## Abstract

CRISPR/Cas9-based transcriptional activation (CRISPRa) has recently emerged as a powerful and scalable technique for systematic over-expression genetic analysis in *Drosophila melanogaster.* We present flySAM, a potent new tool for *in vivo* CRISPRa, which offers a major improvement over existing strategies in terms of effectiveness, scalability, and ease-of-use. flySAM outperforms existing *in vivo* CRISPRa strategies, and approximates phenotypes obtained using traditional Gal4-UAS over-expression. Further, because flySAM typically only requires a single sgRNA, it dramatically improves scalability. We use flySAM to demonstrate multiplexed CRISPRa, which has not been previously shown *in vivo.* In addition, we have simplified the experimental usage of flySAM by creating a single vector encoding both the UAS:Cas9-activator and the sgRNA, allowing for inducible CRISPRa in a single genetic cross. flySAM will thus replace previous CRISPRa strategies as the basis of our growing genome-wide transgenic over-expression resource, TRiP-OE.

## Introduction

A number of techniques have recently been developed for systematic gene over-expression studies based on CRISPR/Cas9 transcriptional activators (CRISPRa) (1). In CRISPRa, nuclease-dead Cas9 (dCas9) is used to guide a transcriptional activator complex to a target gene’s transcriptional start site (TSS) via a short guide RNA (sgRNAs). CRISPRa offers several advantages over traditional techniques for over-expression (reviewed in 2), and represents an important complement to existing genome-wide resources for loss-of-function studies (3). A central challenge now is to adapt CRISPRa for *in vivo* use, to allow for robust, systematic overexpression studies in an organismal context (2, 4-7).

We have recently shown that dCas9-VPR is an effective tool for CRISPRa *in vivo* in *Drosophila* (2, 7). In VPR, dCas9 is fused to a tripartite transcriptional activator domain (VP64-p65-Rta) (8). While VPR successfully activates target genes and generates phenotypes *in vivo*, this system suffers from three important limitations. First, VPR typically requires two sgRNAs per target gene to reliably achieve consistent transcriptional activation (2). This greatly increases the cost and complexity of creating a large-scale resource for *in vivo* CRISPRa, and also doubles the chance of off-target effects. Second, because CRISPRa requires three independent transgenes in a single fly (Gal4, UAS:dCas9-VPR, and sgRNA), the usage of this system is not as straightforward as standard Gal4-UAS based tools which only require a single genetic cross. Third, previous experiments in *Drosophila* cell culture suggest that an alternative CRISPRa technique, synergistic activation mediator (SAM), outperforms VPR in direct comparisons (1). However, previous attempts to express SAM components *in vivo* have failed due to toxicity (2).

In contrast to VPR, SAM involves two separate protein components, dCas9-VP64 and MCP:p65-HSF1, as well as a modified sgRNA containing two MS2 hairpin structures that recruit MCP:p65-HSF1(9). To date, it has not been possible to apply SAM *in vivo* in flies because ubiquitous expression of UAS:MCP-p65-HSF1 is lethal in the absence of any sgRNA (2). In addition, a number of attempts to express “SAM-like” components, including alternative transcriptional activation domains fused to MCP, failed to outperform VPR *in vivo* (2). For these reasons, the first generation of transgenic lines for *in vivo* CRISPRa (the “TRiP-OE” collection) was based on VPR (2).

Here, we present flySAM, a robust, scalable, and simplified strategy for *in vivo* CRISPRa that overcomes previous toxicity issues and unambiguously outperforms VPR in direct comparisons. We show that flySAM using a single sgRNA typically outperforms VPR using two sgRNAs, and that the severity of flySAM phenotypes is comparable to traditional Gal4-UAS over-expression. We use flySAM to demonstrate multiplexed CRISPRa of multiple genes for the first time *in vivo.* Lastly, we have greatly simplified the use of this system by combining flySAM with the sgRNA in a single transgenic vector, allowing for tissue-specific CRISPRa with a single genetic cross. flySAM thus represents a major improvement in the strength, scalability, and ease of use over existing CRISPRa strategies, and will replace VPR as the basis of our growing genome-wide transgenic CRISPRa resource, TRiP-OE.

## Results & Discussion

**Establishing *in vivo* flySAM using the T2A self-cleaving peptide.** Previous attempts to express the SAM component UAS:MCP-p65-HSF1 *in vivo* utilized the *pJFRC7* vector, which is designed for very high protein expression levels, and includes 20XUAS binding sites and a 5’ intervening sequence (IVS) to boost translational efficiency (2). We hypothesized that expressing MCP:p65-HSF1 at lower levels may overcome the lethality observed in previous reports (2). We therefore designed a new vector, flySAM1.0, expressing the SAM components dCas9-VP64 and MCP-p65-HSF1 separated by a T2A self-cleaving peptide, under 10XUAS control (Figure 1A), reasoning that proteins in the second position of a T2A-containing bi-cistronic transcript are typically expressed at lower levels than proteins in the first position (10). We also constructed SAM-compatible sgRNA expression backbone vectors for expressing either a single sgRNA (U6:2-sRNA2.0) or multiple sgRNAs (see Methods).

**Figure 1.**
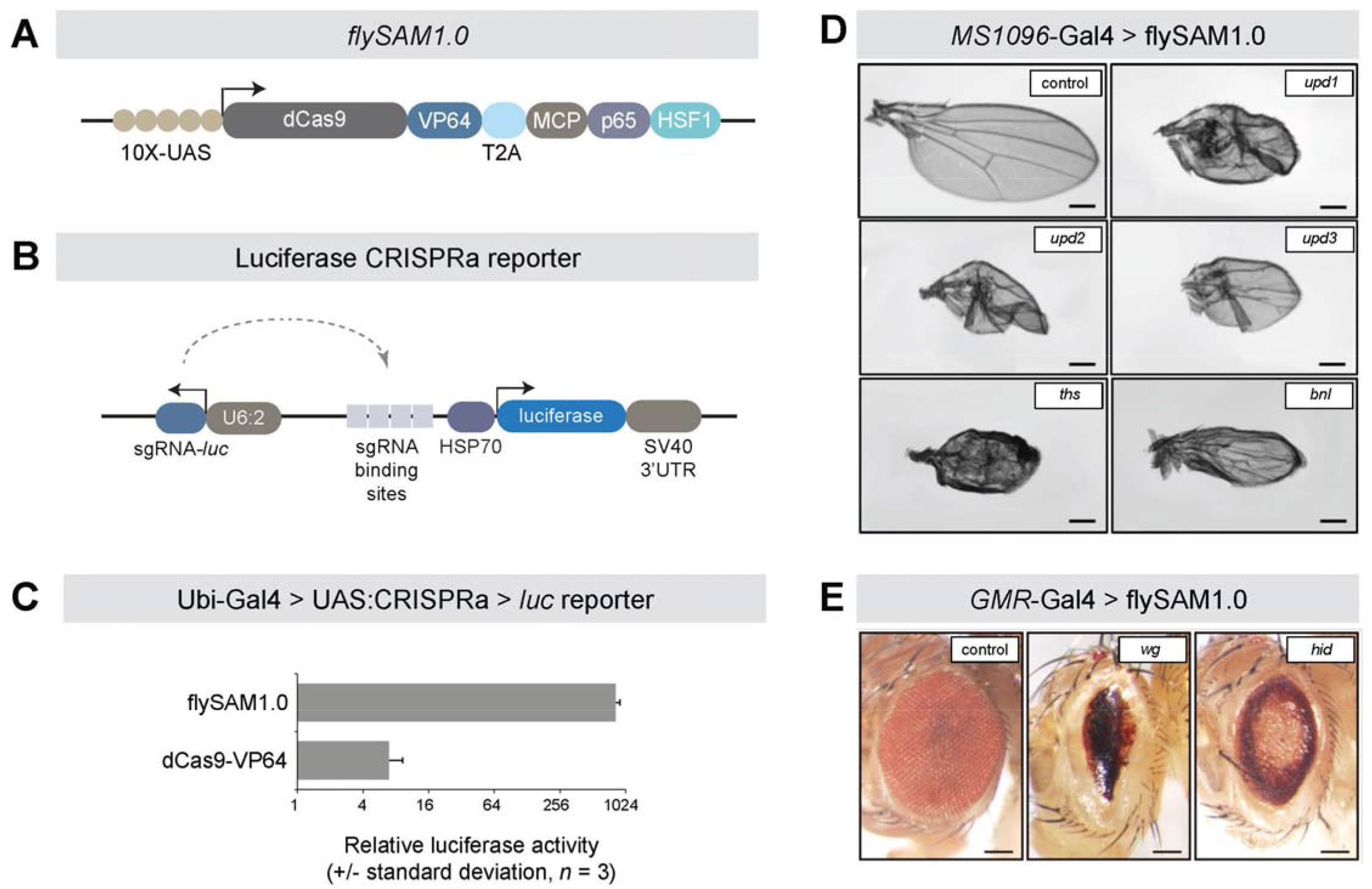
**flySAM1.0 is a potent technique for *in vivo* CRISPRa.** (A) Schematic of the flySAM1.0 construct. The two SAM components are separated by a T2A peptide. Not drawn to scale. (B) Schematic of the luciferase CRISPRa reporter, containing the luciferase coding sequence downstream of four tandem binding sites for an sgRNA that is encoded as a separate transcript. (C) *In vivo* CRISPRa luciferase assay of flySAM1.0 compared to dCas9-VP64, driven by Ubi-Gal4. (D) flySAM1.0 activates a range of endogenous genes *in vivo* in the wing (MS1096-Gal4) and (E) eye (GMR-Gal4), using a single sgRNA. Scale bars = 250 μm.

We tested whether flySAM1.0 can be expressed *in vivo* without toxicity by crossing this line to a ubiquitous Gal4 (actin-Gal4) in the absence of any sgRNA. In contrast to the complete lethality observed for 20XUAS-IVS-UAS:MCP-p65-HSF1(2), we observed normal survival rates, and no visible phenotypes for flySAM1.0 (Figure S1). Similar results were obtained for a second ubiquitous driver, Ubi-Gal4. To further confirm that flySAM1.0 is not toxic *in vivo*, we crossed this line to a panel of tissue-specific Gal4 drivers (hh-Gal4, *MS1096-Gal4, dMef2-*Gal4, *Lpp*-Gal4, *nub*-Gal4). In all cases, we observed normal survival rates, and no visible morphological phenotypes in any of the targeted tissues. We did observe, however, that expression of flySAM1.0 using *tubulin*-Gal4, an additional ubiquitous driver, was lethal, indicating that this construct may be toxic when expressed ubiquitously at high levels.

To test whether flySAM successfully activates target genes *in vivo*, we constructed a luciferase reporter line containing the *luciferase* coding sequence downstream of four tandem sgRNA binding sites, as well as an sgRNA targeting these binding sites, together in one vector (Figure 1B). Using this reporter line, we measured the activity of UAS:flySAM1.0 relative to a UAS:dCas9-VP64 control using Ubi-Gal4, and observed a dramatic increase in luciferase activity, indicating that flySAM1.0 is effective *in vivo* (Figure 1C). We also constructed two additional flySAM-like vectors containing alternative activation domains in place of MCP-p65-HSF1 (flySAM1.1 = UAS:dCas9-VP64-T2A-MCP-p65-Rta and flySAM1.2 = UAS:dCas9-VP64-T2A-MCP-Dorsal-dHSF). However, both of these lines had reduced survival or full lethality when expressed ubiquitously using *actin*-Gal4, and furthermore failed to outperform flySAM using our luciferase reporter assay with Ubi-Gal4 (Figure S1).

Next, we tested whether flySAM1.0 is capable of activating endogenous genes. We generated U6:2-sgRNA2.0 lines targeting a number of genes: the secreted ligands *upd1, upd2, upd3, ths, bnl*, and *wg*, and the pro-apoptotic gene hid. We drove flySAM in the wing (using MS1096-Gal4) or the eye (GMR-Gal4), and observed specific phenotypes for each sgRNA, indicating that flySAM1.0 can trigger physiologically relevant levels of transcriptional activation *in vivo* (Figure 1D,E). We note that these phenotypes were observed using a single sgRNA, while VPR typically requires two sgRNAs for effective CRISPRa (2, 7).

**flySAM outperforms VPR in direct comparisons.** To directly compare flySAM1.0 to VPR *in vivo*, we first used our luciferase reporter system, either containing MS2 hairpins (for flySAM1.0) or a standard sgRNA tail (for VPR). flySAM1.0 led to far higher levels of luciferase activity than VPR (Figure 2A). The superior performance of flySAM is not due solely to MS2 loops in the sgRNA, as the presence of MS2 loops in fact decreased the performance of dCas9-VPR in cell culture (Figure S2).

**Figure 2.**
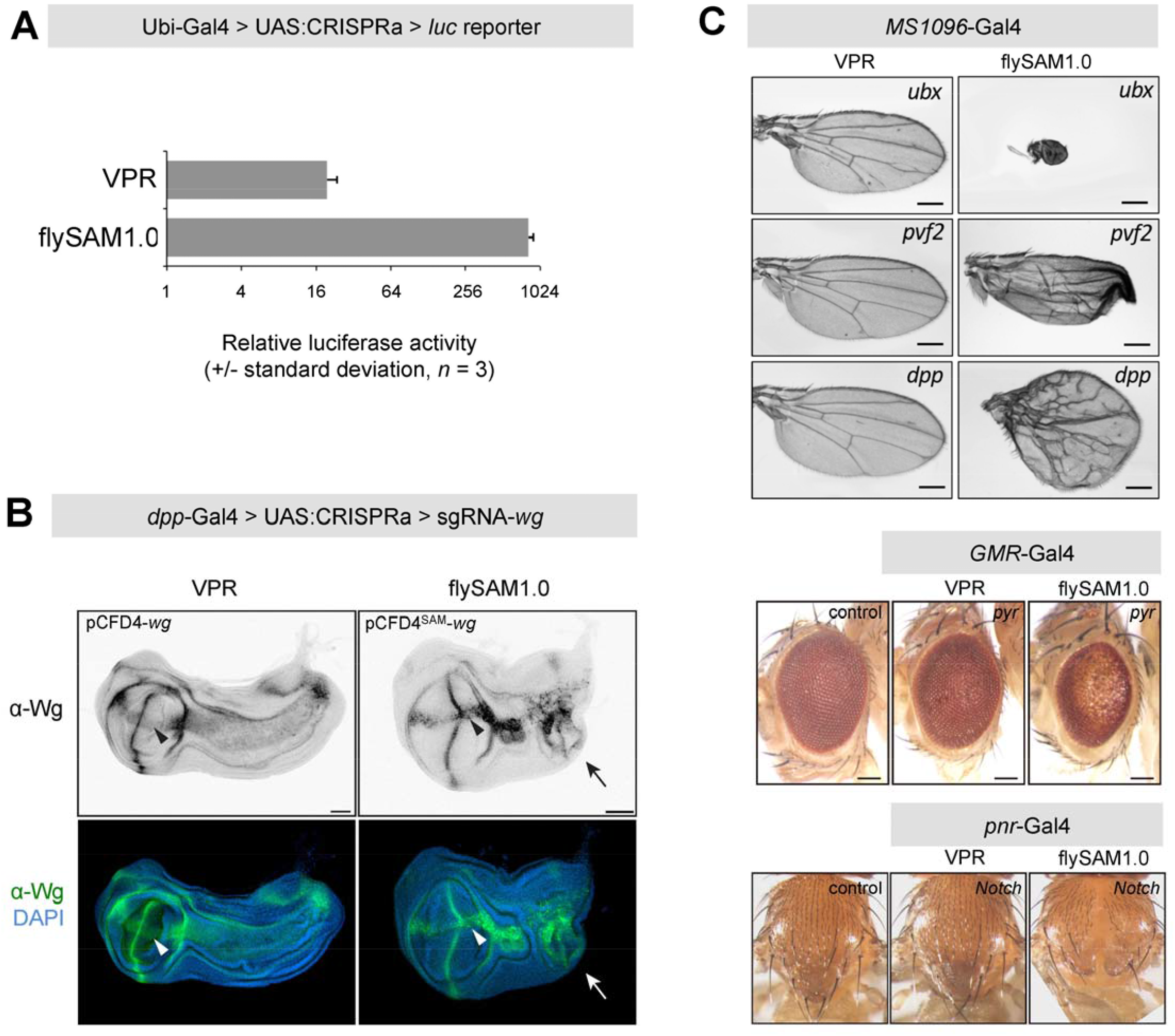
**flySAM1.0 outperforms VPR.** (A) In vivo CRISPRa luciferase assay of flySAM1.0 compared to dCas9-VPR, using Ubi-Gal4. (B) Ectopic activation of wg in the larval wing disc using *flySAM1.0* versus *VPR.* Arrowhead indicates duplicated wing pouch, arrows indicate ectopic Wg expression. Scale bar = 50 μm. (C) Comparison of flySAM1.0 and VPR in the wing, eye, and notum, using identical protospacer sequences were used in all experiments, differentiated only by the presence of MS2 loops in the flySAM experiments. Scale bars = 250 μm.

To compare the ability of VPR and SAM to activate endogenous genes *in vivo*, we utilized an established system for comparing CRISPRa activity, wherein dpp-Gal4 is used to activate *wg* in an ectopic domain of the larval wing imaginal disc (2, 7). In these experiments, two sgRNAs were used, which were identical in protospacer sequence, and differ only in the presence of MS2 loops in the sgRNA tail. While both systems activated ectopic Wg expression, flySAM1.0 induced a duplication of the wing pouch, complete with a secondary dorso-ventral (D-V) axis, and visibly higher levels of ectopic Wg expression (Figure 2B). A fully duplicated wing pouch is reminiscent of the phenotype observed using *dpp-Gal4* > *UAS:wg* (11), and is never observed using VPR. The phenotype observed with flySAM1.0 is thus unambiguously stronger than VPR.

Previous experiments comparing VPR and SAM in cell culture have shown that, across multiple individual tests of different genes, there are some cases where VPR can outperform SAM (1). We therefore tested an additional five genes *(ubx, Pvf2, dpp, pyr, N;* all using single sgRNAs) in different tissues (wing - MS1096-Gal4; eye - GMR-Gal4; notum - pnr-Gal4), and found that in all cases flySAM generated stronger phenotypes than VPR (Figure 2C). Thus, we conclude that, given any individual sgRNA, flySAM outperforms VPR *in vivo.*

**flySAM induces phenotypes resemble UAS:cDNA overexpression phenotypes.** Traditionally, the most widely used technique for *in vivo* overexpression in *Drosophila* is cDNA overexpression using the Gal4-UAS system (12, 13). To directly compare flySAM to cDNA overexpression, we used U6:2-sgRNA2.0 lines for five genes for which UAS overexpression lines are available (*wg, hh, ci, dpp*, and *Ras85D*). Driving these constructs in the wing (MS1096-Gal4) or eye (eyeless-Gal4), we observed comparable phenotypes in all cases (Figure 3). These results indicate that flySAM1.0 recapitulates established overexpression phenotypes comparable in strength to Gal4-UAS overexpression, using only a single sgRNA.

**Figure 3.**
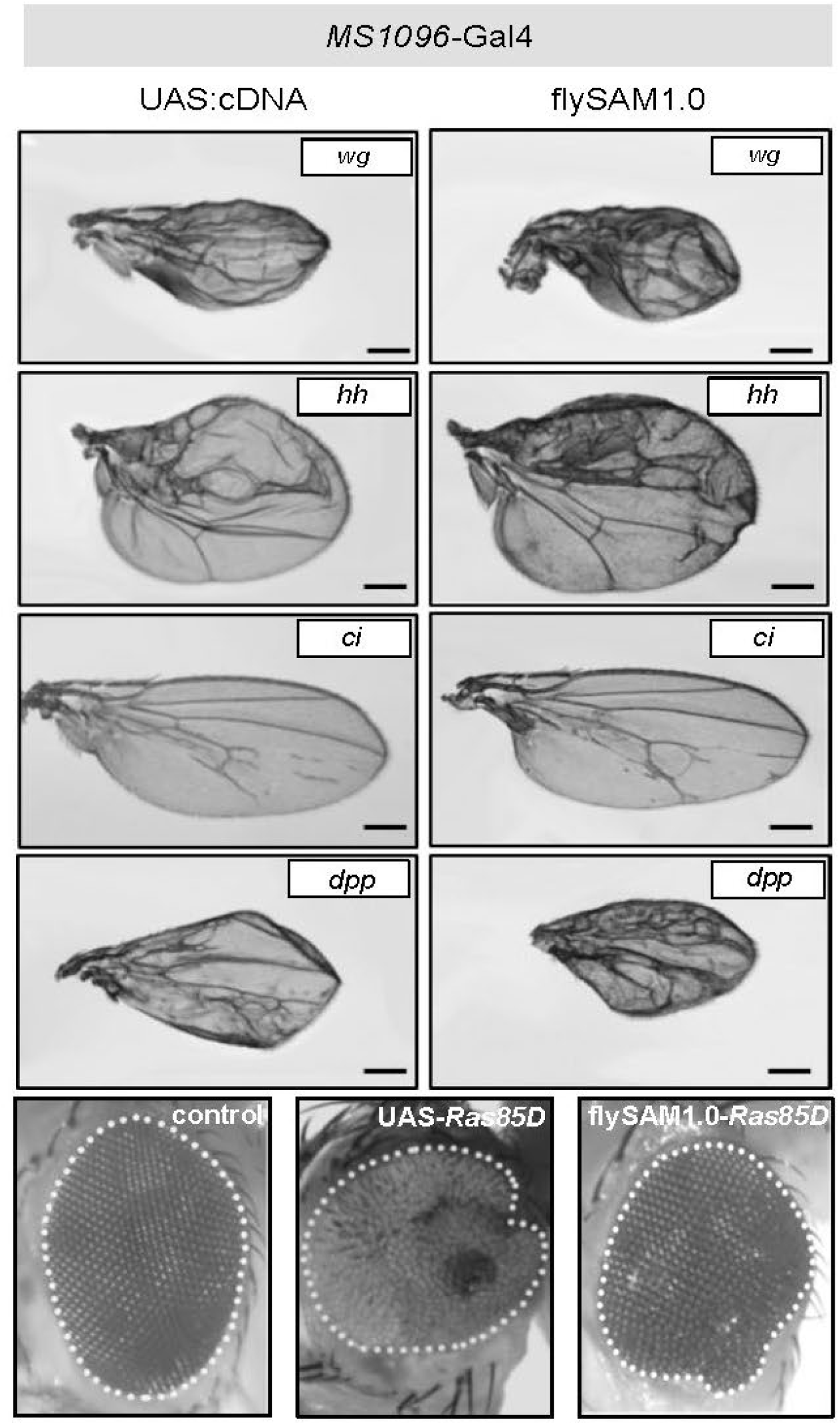
**flySAM1.0 phenotypes recapitulate Gal4-UAS over-expression phenotypes.** The phenotypes observed using flySAM1.0 (with one sgRNA) resemble those obtained using *Gal4* > *UAS-cDNA.* Wing phenotypes = MS1096-Gal4; eye phenotype = *eyeless*-Gal4. Scale bars = 250 μm.

**flySAM allows for multiplexed CRISPRa.** We tested whether it is possible to activate multiple genes simultaneously by co-expressing multiple sgRNAs in a single fly. We first activated *ci* and hh, both singly and together, in the wing using MS1096-Gal4, using a single sgRNA vector that expresses either one or both sgRNAs. When *ci* and *hh* were activated simultaneously, both individual phenotypes were observed in the wing (Figure 4A), indicating successful multiplexed CRISPRa. Next, we tested eye-specific CRISPRa of genes from the Hippo pathway using ey-Gal4. Individual overexpression of *wts, mats*, or *hpo* (2-3 sgRNAs per gene) produced relatively mild small-eye phenotypes, while multiplexed CRISPRa for *wts* + *mats* or *mats* + *hpo* showed strong enhancement of this phenotype (Figure 4B). Similarly, while individual CRISPRa of the tumor suppressors *Tsc1* and *Tsc2* had minimal effects, *Tsc1* + *Tsc2* showed a strong growth-suppressing genetic interaction (Figure 4C). To our knowledge, these results provide the first demonstration of multiplexed *in vivo* CRISPRa.

**Figure 4.**
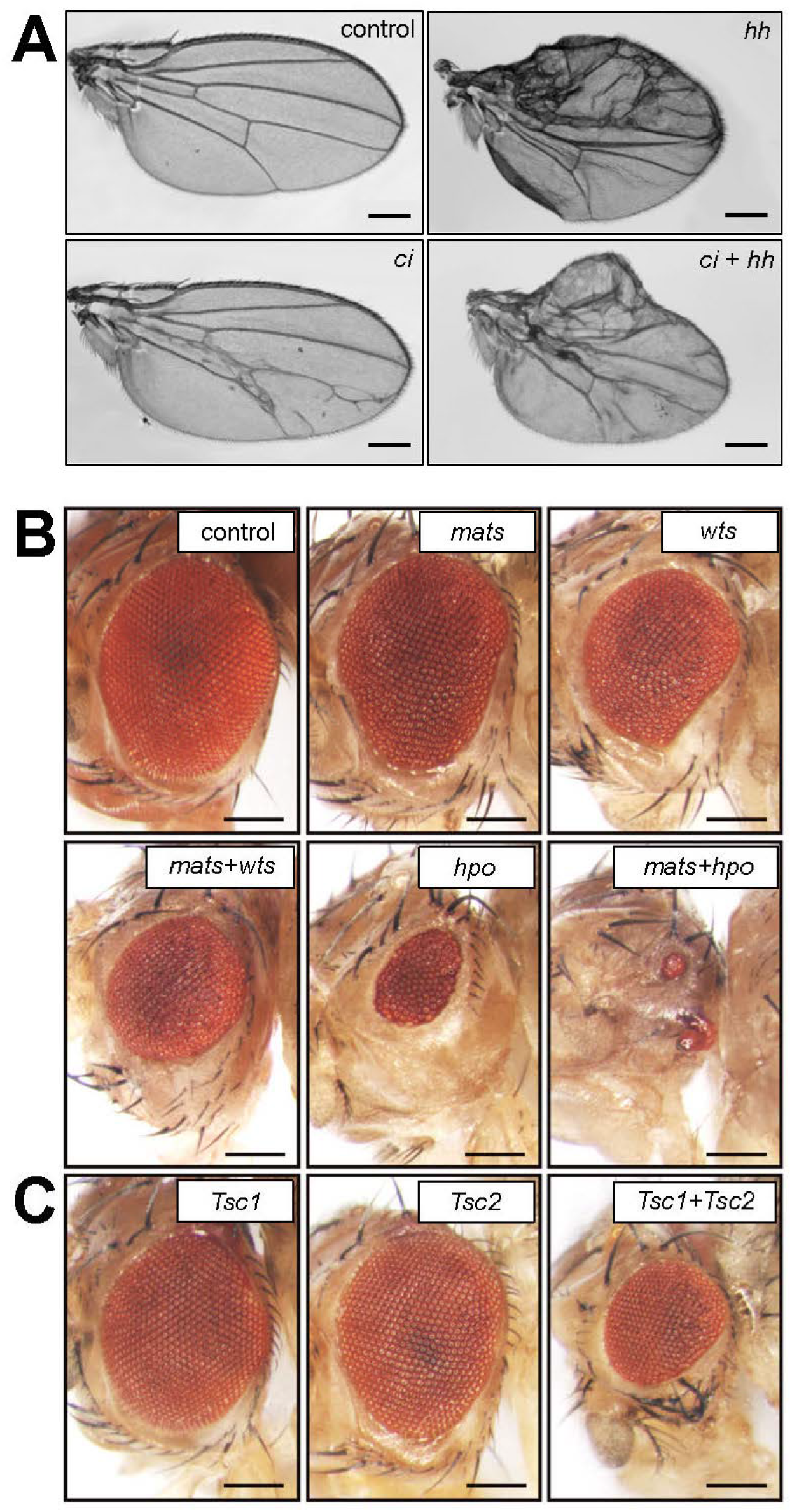
***in vivo* multiplexed CRISPRa using flySAM.** (A) Multiplexed transcriptional activation of *ci* and *hh* in the wing (using MS1096-Gal4) produces both of the individual phenotypes. (B) Multiplexed transcriptional activation of Hippo pathway components in the eye (using eyeless-Gal4). (C) Simultaneous CRISPRa of *Tsc1* + *Tsc2* in the eye. In panels B and C, note that 2-3 sgRNAs per target gene were used.

**flySAM2.0 - a single vector containing UAS:flySAM and sgRNA for simplified experimental set-up.** A major technical drawback of both the VPR system and the flySAM system described above is that they involve three separate transgenic elements: a Gal4, a UAS-CRISPRa construct, and an sgRNA construct. Because of this, it requires the creation of a compound fly stock containing two of the components before crossing this compound stock to a line expressing the third component. In our previous work with VPR, we attempted to overcome this technical bottleneck by generating a publicly available collection of over 30 Gal4 + UAS:VPR compound stocks containing widely-used Gal4 lines combined with UAS:VPR (2). However, utilizing any additional Gal4 requires the generation of a compound Gal4 + UAS stock before performing CRISPRa experiments, which takes several weeks.

To simplify *in vivo* CRISPRa, we created a single vector containing both UAS:flySAM and a sgRNA, termed flySAM2.0 (Figure 5A), reasoning that flySAM2.0 lines could simply be crossed to a Gal4 line to achieve tissue-specific CRISPRa in the offspring. We also included *gypsy* elements flanking the CRISPRa components, in order to insulate the CRISPRa components from surrounding regulatory features that could dampen their expression (14).

**Figure 5.**
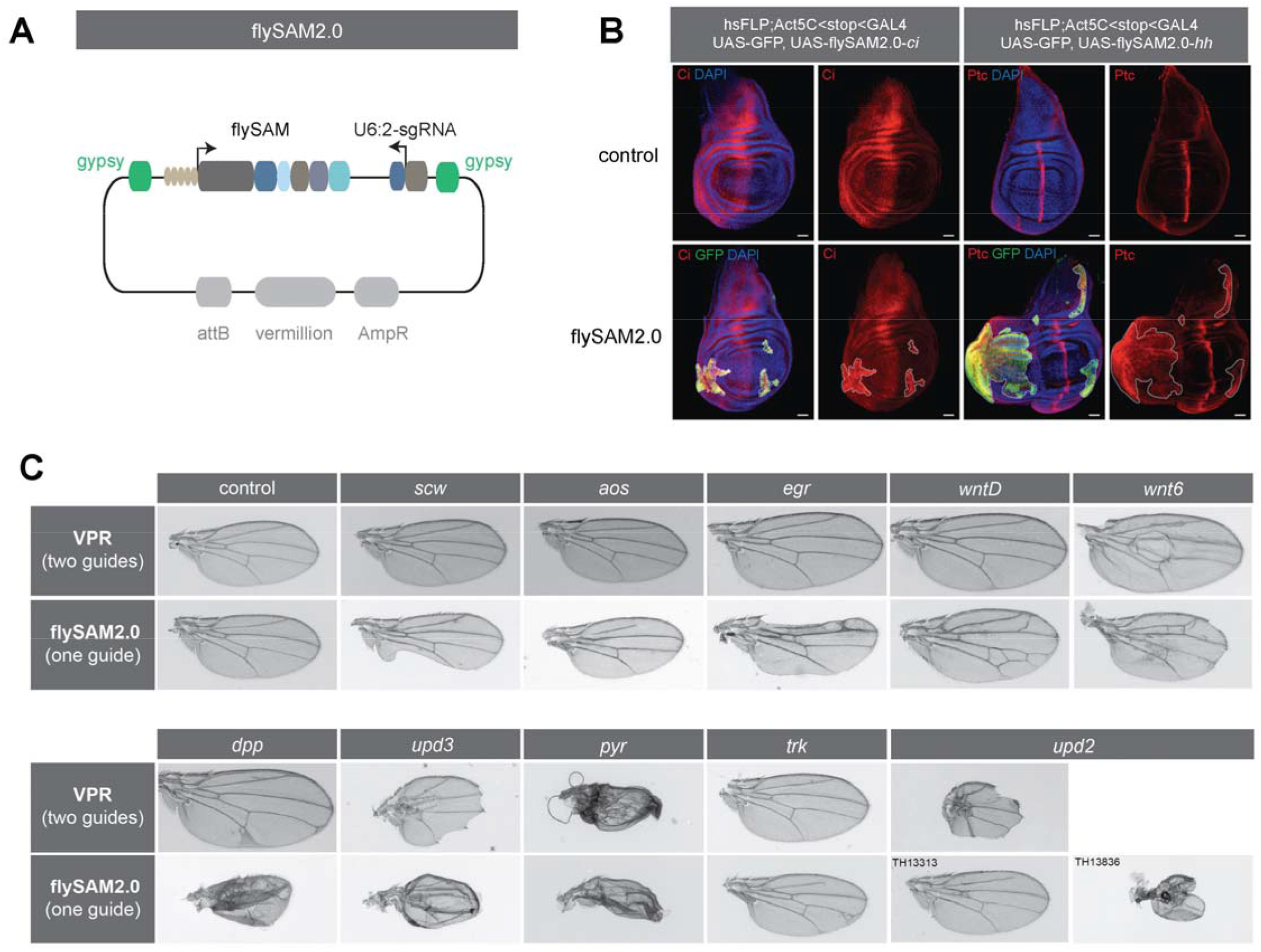
**flySAM2.0 allows for inducible CRISPRa with a single genetic cross.** (A) Diagram of flySAM2.0, which contains both the UAS:flySAM and the sgRNA in a single plasmid, between *gypsy* insulators. (B) Clonal CRISPRa using *FLPout Gal4* > *flySAM2.0* for *ci* and *hh* in L3 larval wing discs; GFP+ clones are indicated with dashed white lines. Scale bar = 50 μm. (C) Comparison of flySAM2.0 (one sgRNA per target gene) to an existing VPR collection (using two sgRNAs per target gene). See text for details.

To verify that flySAM2.0 functions as intended, we generated two flySAM2.0 lines targeting either *hh* or *ci*, crossed both lines to a FLPout-Gal4 stock *(hsFlp; actin-FRT-STOP-FRT-Gal4; UAS-GFP)*, and examined CRISPRa clones in the larval wing disc using antibody staining for Ci or the *hh* target gene Ptc. In both cases we detected strong, specific activation in GFP+ clones, indicating successful CRISRPa in a single genetic cross (Figure 5B). These experiments also confirm previous observations that CRISPRa is highly specific, as we never observed CRISPRa outside of Gal4+ clones (2).

Having established that flySAM outperforms VPR for any given sgRNA, we tested whether flySAM2.0 (with a single sgRNA) outperforms the existing collection of VPR sgRNA stocks (two sgRNAs per target gene). To do so, we compared CRISPRa wing phenotypes *(nub*-Gal4) using independent lines generated for each strategy, and which thus differ in their protospacer sequence (Table S2). We first focused on a set of target genes for which existing VPR had previously failed to generate a phenotype (*scw, aos, egr, WntD*, and *trk*), or had been relatively weak (dpp). For all six of these genes, flySAM2.0 generated strong, specific phenotypes that were not observed using VPR (Figure 5C). We also tested a number of genes for which VPR did show phenotypes (*upd2, upd3, pyr*, and *Wnt6*). For *upd3, pyr*, and *Wnt6*, flySAM2.0 and VPR produced comparable phenotypes, whereas for *upd2*, VPR produced a stronger phenotype than flySAM2.0 (Figure 5D). We attribute this result to the fact that because flySAM2.0 only uses a single sgRNA, this is likely a case where that single sgRNA is sub-optimal. Accordingly, a second independent flySAM2.0 line targeting *upd2* generates a strong phenotype in the wing (Figure 5D). Thus, taken together our results demonstrate that flySAM2.0 with a single sgRNA outperforms VPR with two sgRNAs in the large majority of cases. Going forward, we refer to flySAM2.0 as simply “flySAM” for simplicity.

We note that the flySAM construct is substantially larger than a typical sgRNA construct (15 kb vs. 7 kb), which is likely to reduce the efficiency of recovering transgenic lines. To address this concern, we directly compared the recovery of transformants from a pooled injection of 16 U6:2-sRNA2.0 constructs versus 16 flySAM constructs. While the total number of transformants recovered was lower for flySAM (flySAM = 26 transformants from 117 surviving injected embryos; U6:2-sgRNA2.0 = 83 transformants from 108 surviving injected embryos; Table S3), we were still able to recover 10 of the 16 constructs for flySAM, compared to 13 of the 16 constructs for U6:2-sgRNA2.0 (Table S3). Given the greatly improved downstream usefulness of flySAM constructs, and given that we can still recover transformants at a reasonable rate, we have begun large-scale production of flySAM constructs, towards a long-term goal of generating a genome-wide *in vivo* CRISPRa resource.

In summary, flySAM represents a major improvement over existing techniques for *in vivo* CRISPRa in terms of effectiveness, scalability, and ease-of-use. Regarding effectiveness, flySAM outperforms VPR in direct comparison holding sgRNA sequence constant (Figure 2), and in fact flySAM using a single sgRNA outperforms VPR using two sgRNAs in the large majority of cases (Figure 5). Regarding scalability, flySAM dramatically decreases the cost and time required to generate transgenic constructs. Specifically, while double-sgRNA constructs require PCR, gel purification, and Gibson cloning (15), single sgRNA constructs only require ligation of annealed oligos into a plasmid backbone. Finally, by combining the SAM components and sgRNA in a single vector, flySAM provides simplicity, allowing researchers to perform tissue-specific CRISPRa in a single genetic cross.

Given these improvements, we anticipate that flySAM will be widely useful to the *Drosophila* community for over-expressing specific genes in an inducible, tissue-specific manner, as well as performing screens. We have begun transitioning our growing collection of transgenic TRiP-OE CRISPRa stocks from VPR to flySAM. Details of this collection are available at https://fgr.hms.harvard.edu/fly-in-vivo-crispr-cas.

## Methods

### *Drosophila* stocks

All Drosophila stocks were maintained at 25°C with 60% humidity on standard cornmeal/sugar/agar media unless otherwise specified. CRISPRa lines (Table S1) and sgRNA lines (Table S2), were obtained from the Tsinhua Fly Center (THFC) or the Bloomington Drosophila Stock Center (BDSC). Additional fly stocks were as follows: *UAS-wg* (a gift from Dr. Ting Xie), *UAS-hh* (TH13350.S), *UAS-ci* (TH11173.N), *UAS-dpp* (THJ0143, from Dr. Jose C. Pastor-Pareja), *UAS-Ras85D* (THJ0149, from Dr. Jose C. Pastor-Pareja), *y,hsFLP; act5C* < *y* +< *GAL4,UAS-GFP/CyO* (a gift from Dr. Hong Luo), *MS1096-GAL4* (Bloomington 8860), *Ubi-GAL4* (16), *ey-Gal4* (Bloomington 5535), *dpp-Gal4* (Bloomington 7007), *ptc-Gal4* (Bloomington 2017) *GMR-Gal4* (Bloomington 8605), dMef2-Gal4 (17), and Lpp-Gal4 (18). nub-Gal4, hh-Gal4, actin-Gal4, and *tubulin-Gal4* are Perrimon Lab stocks. UAS-hh and UAS-ci were created by first amplifying the coding sequences of these genes from Drosophila genomic DNA, and then cloned into a pVALIUM vector (19) with EcoRI and NheI restriction digests, using Hieff Clone™ one step cloning kit (Yeasen).

### Transgene constructs and production of transgenic flies

The flySAM1.0 vector was constructed on the basis of VALIUM20 (19). Fragments were stitched together using the sequence- and ligation-independent cloning (SLIC) (20). dCas9 was amplified from a nos-Cas9 vector (21) and two mutations in the RuvC domain (D10A) and HNH domain (H840A) were introduced separately. Activation domains were amplified from plasmid or genomic DNA by using High-Fidelity DNA Polymerase and inserted into the C terminus of dCas9. VP64, T2A, and MCP were synthesized by GENEWIZ (Suzhou, China). p65 was cloned from mouse cDNA for flySAM1.0 or Addgene 63798 (human) for flySAM2.0. HSF was amplified from Addgene 61426. Rta was amplified from Addgene 63798, Dorsal and dHSF were cloned from *Drosophila* cDNA.

U6:2-sgRNA2.0 vector was cloned by replacing sgRNA1.0 scaffold in U6b-sgRNA-short (21) with the sgRNA2.0 scaffold (9). The flySAM2.0 vector was made by inserting U6B-sgRNA2.0 fragment digested with NheI and SpeI from U6B-sgRNA2.0 vector into flySAM1.0 vector.

We used two separate strategies for expressing multiple sgRNAs from a single plasmid. For dpp-Gal4 > CRISPRa experiments in larval wing disc, we used pCFD4^SAM^ (2), which expresses two sgRNAs from U6:1 and U6:3, respectively. For all other multiplex experiments, we combined multiple U6:2-sgRNA fragments in a single plasmid as follows: first, we cloned individual sgRNAs into the U6:2-sgRNA2.0 vector. One such U6:2-sgRNA2.0 vector was linearized by digesting with SpeI or NheI, while additional vectors were digested with both SpeI and NheI to isolate the fragment containing a U6b-sgRNA2.0 (∼1000bp). The resulting products were gel purified (AxyPrep DNA Gel Exstaction Kit) and ligated together.

To build the luciferase CRISRa reporter, we cloned firefly luciferase downstream of the hsp70 basal promoter, and upstream of an SV40 polyA tail. We inserted a synthesized sgRNA target site (sgRNA = CCTACAGCACGTCGCCGGCG) immediately upstream of the hsp70 promoter. This luciferase reporter fragment was inserted into U6:2-sgRNA2.0 vector using restriction digest cloning to generate sgRNA2.0 luciferase reporter.

Transgenic fly lines were generated by injecting the constructs into *y sc v nanos-integrase; attP2* or *y sc v nanos-integrase; attP40* embryos following standard procedures (19).

### FLP-out induced clonal analysis and Immunostaining

Clones of CRISPRa cells in the wing disc were generated by FLP/FRT-mediated recombination. To generate CRISPRa clones, the flySAM2.0-ci *and* flySAM2.0-hh *lines* were crossed with *y, hs-FLP; act5C<STOP<Gal4, UAS-GFP/CyO* flies and maintained at 25°C. 1^st^ instar larvae were heat shocked at 37°C for 1 hour. Wing discs were dissected from late 3^rd^ instar larvae, stained using standard antibody staining protocol, and imaged using a Zeiss LSM780 confocal.The following primary antibodies were used: anti-Wg antibody (4D4, DSHB, 1:100) rat anti-ci antibody 2A1-c (DSHB, 1:10), mouse monoclonal anti-ptc antibody ptc-c (DSHB, 1:50), and rabbit polyclonal anti-GFP (Abcamab290, 1:2000). Various secondary antibodies (Jackson ImmunoResearch Laboratories) conjugated with FITC or TRITC were used at 1:300.

**Luciferase assay.** Luciferase activity was measured using the Steady-Glo Luciferase Assay Kit (Promega, E2520) as previously described (22). Briefly, three 7-day-old adult female flies were collected separately in 100μl or 50μl Glo Lysis Buffer (Promega, E266A) for each sample; five independent samples were used for each luciferase assay. Samples were homogenized and then centrifuged for 15min at 20,000rcf, 4°C. 55μl or 35μl of supernatant were transferred to a 96 well solid white microplate and mixed with the same volume of Steady-Glo reagent. After incubation in the dark for 20 min, luminescence was measured on a spectral scanning multimode reader (Thermo Scientific, Varioskan^®^ Flash).

**Comparison of flySAM and VPR.** The sgRNAs targeting *wg* (in either pCFD4 or pCFD4^SAM^) have been previously described (2). These sgRNA-wg lines were crossed to *w; dCas9-VPR; dpp-Gal4* / TM6b or *w; flySAM1.0; dpp-Gal4* / TM6b, respectively, and maintained at 27°C. Wing discs were dissected from non-Tubby L3 larvae, fixed, stained, and imaged as described above..

To directly compare flySAM2.0 collection (one sgRNA, one transgene) to dCas9-VPR (two sgRNAs, two transgenes), a published collection of double-sgRNA lines for VPR (2) was compared to a newly generated collection of single-sgRNA lines built in the flySAM2.0 backbone targeting the same genes. VPR lines were crossed to *w; nubbin-Gal4; dCas9-VPR /SM5, TM6b*, and flySAM2.0 lines were crossed to w;; *nubbin-Gal4*, and all crosses were maintained at 27°C. Adult wings (10-20 per sex, per genotype) were mounted in Permount (Fisher) and analyzed on a Zeiss Axioskop 2 using brightfield optics.

**Cell culture and qPCR.** Cell culture experiments and quantitative PCR were performed as described (2, 7), with primers described in (1).

## Acknowledgements

We thank Ryan Colbeth and Rich Binari for expertise and assistance with fly work. This work was supported by NIH Grants R01GM084947 and R24OD021997 (N.P. laboratory). B.E.-C. was funded through the NIH’s Ruth L. Kirschstein National Research Service Award F32GM113395 from the NIH General Medical Sciences Division. The J.-Q.N. laboratory This work was supported by the National Key Technology Research and Development Program of the Ministry of Science and Technology of the People’s Republic of China (2015BAI09B03, 2016YFE0113700), and the National Natural Science Foundation of China (31571320) N.P. is an investigator of the HHMI.

